# Proposed model of the Dictyostelium cAMP receptors bound to cAMP

**DOI:** 10.1101/694810

**Authors:** Jack Calum Greenhalgh, Aneesh Chandran, Matthew Thomas Harper, Graham Ladds, Taufiq Rahman

## Abstract

3’,5’-cyclic adenosine monophosphate (cAMP) is well known as a ubiquitous intracellular messenger regulating a diverse array of cellular processes. However, for a group of social amoebae or Dictyostelia undergoing starvation, intracellular cAMP is secreted in a pulsatile manner to their exterior. This then uniquely acts as a first messenger, triggering aggregation of the starving amoebae followed by their developmental progression towards multicellular fruiting bodies formation. Such developmental signalling for extracellularly-acting cAMP is well studied in the popular dictyostelid, *Dictyostelium discoideum*, and is mediated by a distinct family (‘class E’) of G protein-coupled receptors (GPCRs) collectively designated as the cAMP receptors (cARs). Whilst the biochemical aspects of these receptors are well characterised, little is known about their overall 3D architecture and structural basis for cAMP recognition and subtype-dependent changes in binding affinity. Using a ligand docking-guided homology modelling approach, we hereby present for the first time, plausible models of active forms of the cARs from *D. discoideum*. Our models highlight some structural features that may underlie the differential affinities of cAR isoforms for cAMP binding and also suggest few residues that may play important roles for the activation mechanism of this GPCR family.

## 1. Introduction

G protein-coupled receptors (GPCRs) represent a large family of evolutionarily-conserved, transmembrane signalling proteins that recognise various changes in the extracellular milieu and transduce them into the cell interior, thereby enabling the generation of appropriate cellular responses. An important example of the latter is chemotaxis, which involves directed migration of a cell along the gradient of extracellular chemical cues [1]. This is an essential process for the development and homeostasis of both unicellular and multicellular organisms and is implicated in many pathophysiological conditions. Much of our current understanding of GPCR-mediated chemotaxis signaling is derived from studies on the social amoeba, *Dictyostelium discoideum*, and mammalian neutrophils [2]. The chemotaxis of *D. discoideum* during starvation is by now a well-established paradigm where cAMP uniquely plays a key role as an ‘extracellular’ (i.e. first) messenger, in stark contrast with its canonical role as a major intracellular (i.e. second) messenger. In its natural habitat (e.g. deciduous forest soil and decomposing leaves), *D. discoideum* feeds on bacteria and yeast and grows individually. When the food source is exhausted, the amoebae aggregate to form a multicellular slug that then differentiates into a fruiting body. Such starvation-induced aggregation of individual *D. discoideum* is driven by propagating waves of extracellular cAMP serving as the chemoattractant. cAMP is first released in a pulsatile manner by a few starving cells that typically lie in the middle of the aggregation centre. The neighbouring cells then recognise the signal via cell surface cAMP receptors (cARs), activate the aggregation-specific adenylyl cyclase (ACA) and produce and secrete additional cAMP (‘cAMP relay’). The latter ultimately results in outward propagating waves of cAMP, which direct inward movement of the cells towards the aggregation centre. cAMP binding to the cARs also triggers receptor desensitization, which results in a cessation of ACA activation and downregulation of cAMP production. A highly specific secreted cAMP phosphodiesterase (PdsA) causes the extracellular cAMP levels to fall. When cAMP levels drop, the cells de-adapt with cARs regaining sensitivity and another wave of cAMP stimulation occurs. In addition to chemotaxis, the repetitive activation of cARs is also a major regulator controlling the expression of developmentally regulated and cell-type specific genes [2, 3].

As described above, the key proteins that enable the chemoattractant role of cAMP are cARs, which are now categorised as members of the class E sub-family of GPCRs [4]. As the chemoattractant, cAMP binds to surface receptors that are encoded by four genes, *cAR1*, *cAR2*, *cAR3* and *cAR4*, each of which is transiently expressed at different stages of *D. discoideum* development during starvation. All of the encoded proteins (cAR1-cAR4) appear to be coupled largely to similar signal-transduction pathways but differ in the temporal and spatial aspects of their expression pattern and more importantly in their affinities for binding cAMP [5]. The cAMP-binding affinities of these receptors differ significantly but are correlated with the stage of their expression. cARl appears during the early aggregation stage, whereas cAR3 is maximal during the mound stage. These are followed by cAR2 and cAR4, expressed exclusively in pre-stalk cells in the slug and culmination stages, respectively [5, 6]. Molecular phylogenetic analyses and subsequent functional studies have shown cAR1 orthologues to be present in most other members of the Dicotyostelia but not all of them use extracellular cAMP (and thus the cAR1-like proteins) to drive aggregation unlike those belonging to the subgroup represented by *D. discoideum*. For the other three subgroups, cAR1 orthologues are expressed after aggregation and oscillatory extracellular cAMP signalling mediated through these receptors deem indispensable for a more conserved role such as fruiting body morphogenesis [7].

The highest affinity cAMP receptor encoding *cAR1* was first cloned by Peter Devreotes and colleagues. Initial hydropathy analysis predicted seven transmembrane (7-TM) topology and revealed significant sequence homology with few GPCR sequences known during that time [8]. Subsequent studies by the same group led to the identification of genes and cloning of sequences of the other three *cARs* that encode cAR2, cAR3 and cAR4 proteins [5, 9]. The cAR isoforms share approximately 60% sequence identity within their predicted transmembrane and loop regions but seem to differ more in their C-terminal regions [9, 10]. It is also well established that downstream of cAMP binding to all cARs, βγ subunit dissociates from the Gα2 and activates ACA to generate intracellular cAMP but phospholipase C and guanylyl cyclase are also activated [5, 6].

cAR1 is the primary cAR isoform responsible for aggregation since *D. discoideum* strain lacking *cAR1* fail to initiate cAMP signalling and fail to aggregate [8, 11]. Given the riche palette of available 3D structures of GPCRs hitherto available in the protein data bank (PDB) and given the indispensability of cAR1 for the age-old paradigm of the starvation-induced *D. discoideum* aggregation, we sought out to model the 3D structure of agonist-bound cAR1 protein first, using a systematic and comprehensive approach guided by plausible binding mode of cAMP. Based on our learning from this cAR1 modelling, we then built homology models of the other three cAR proteins in complex with cAMP. There were several reasons behind why we undertook such modelling task. First of all, to date there is no published 3D structure of any member belonging to the class E family of GPCRs. There also appears to be paucity of comprehensive analysis on this GPCR family members from the structural point of view, especially when substantial improvement and success rate in elucidating 3D structures of GPCRs using X-ray and cryo-EM have been achieved in last 10 years. The cARs seem to lack any sequence or motif compatible with the canonical cyclic nucleotide binding (CNB) domain [12] present in other eukaryotic and prokaryotic cAMP binding proteins, all of which are intracellular including notably PKA, cyclic nucleotide gated (CNG) ion channels, exchange protein directly activated by cAMP (Epacs) and the catabolite activator protein (CAP). We therefore reckoned that our modelling approach may reveal a novel cAMP binding motif and enable us explain the well-established differences in cAMP-binding affinities across cAR isoforms from a structural point of view.

## 2. Methods

### 2.1 Homology modelling

To find the best templates, the sequences of cAR1 (uniprot id: P13773) was first screened against the protein data bank (pdb) using the domain enhanced lookup time accelerated BLAST (DELTA-BLAST) [13]. Sequence alignments of cAR1 and that of the chosen templates were performed using FUGUE (http://mizuguchilab.org/fugue/) [14] and the resultant alignment was used to build 100 homology models per each chosen GPCR template using Modeller 9.23 (http://salilab.org/modeller/). The best models were chosen based on the normalized discrete optimized protein energy (z-DOPE) score implemented within Modeller [15]. For further structure refinement, each of the selected homology models was then embedded in a pre-equilibrated POPC (1- palmitoyl-2-oleoyl phosphatidylcholine) membrane model and subjected to 5ns molecular dynamics (MD) simulation protocol using Gromacs 4.5 and OPLSAA forcefield as per published protocol [16]. 3D quality of the models were evaluated through MolProbity server (http://molprobity.biochem.duke.edu/)[17].

Modelling of cAR2 (uniprot id: P P34907), cAR3 (uniprot id: P35352) and cAR4 (uniprot id: Q9TX43) was performed using the best model of cAR1. The methods of homology modelling, model selection and refinement and 3D quality assessment were essentially the same as for cAR1 mentioned above.

### 2.2 Docking and prediction of binding fee energy

AutoDock 4.2 (http://autodock.scripps.edu/) [18] and GOLD [19] suite version 5.3 (CCDC, Cambridge, UK) were used for blind and focused docking, respectively. For each model, 10-independent docking runs were performed. For selected docked complexes, Molecular Mechanics with Generalized Born Surface Area (MM/GBSA) method as implemented in the Prime 3.0 program in the Schrödinger suite was used to estimate relative free energies (ΔG_bind_) of cAMP binding according to the following equation:

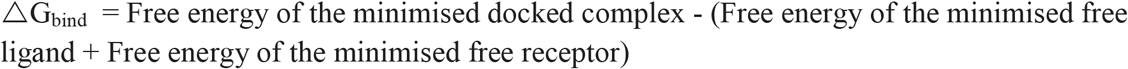

Residues within 5.0 Å of the ligand were allowed to relax during the MM minimisation of the complex, keeping the rest of the structure fixed. We employed the variable dielectric solvent model VSGB 2.0[20], which includes empirical corrections for modelling directionality of hydrogen-bond and π-stacking interactions. This approach has been reported to provide reliable estimates of binding free energies for a wide range of protein–ligand complexes in previous studies [21].

For docking, 3D structure of cAMP was obtained from PubChem (CID: 6076) whilst two analogues of cAMP known to have much reduced affinities for cARs [22–24] were drawn in MarvinSketch (ChemAxon Ltd.) (Figure S1) and subsequently energy minimised using MMFF94 force field using Open Babel 2.4.0[25].

### 2.3 Evolutionary conservation analysis

Analysis and mapping of amino acid conservation onto selected cAR1 models were performed using the Consurf 2016 server (http://consurf.tau.ac.il/2016/) [26]. Both the amino acid sequence (residues 1-268) as well as selected homology models were used as inputs.

### 2.4 Analysis and visualisation of docked complexes

All images of the docked complex were generated using UCSF Chimera 1.14 (https://www.cgl.ucsf.edu/chimera/). Ligand interaction diagrams were generated using PoseView™ [27]. According to the latter, the black dashed lines indicate hydrogen bonds, salt bridges, and metal interactions; green solid line show hydrophobic interactions and green dashed lines show π-π and π-cation interactions.

## 3. Results and discussion

### 3.1 Identification of suitable templates for modelling cAR1

Screening cAR1 sequence against the PDB using DELTA-BLAST (dated: 15-12-2019) using the default parameters (including the statistical significance threshold, E of 0.05) led to >150 hits, all of which are published structures of GPCRs. Visual inspection of the alignments indicated that the C terminus of cAR1 containing 124 amino acids were not aligned with any of the identified hits. We therefore repeated the abovementioned template search excluding this segment of last 124 amino acids and this time we also used a greater statistical threshold (E = 0.001). This produced a hit list of 66 GPCR structures that had significant homology in amino acid sequences with that of cAR1 used as query. It was intriguing to note that the top 14 hits were members from the class B GPCR family. Among other hits as potential templates, there were 8 members of the class F/Frizzled family of GPCRs and the rest were from the class A family of GPCRs (supplementary file). Since a major aim of the present study was to model an agonist (cAMP)-bound and active structure of cAR1, we decided to use active GPCR structures as templates, selected from the individual classes. The first template was the chain R of 4.1Å structure of calcitonin receptor (CTR) -heterotrimeric Gs protein complex solved through cryo-electron microscopy (cryo-EM) (PDB id: 5UZ7) [28]. However, due to better resolution, we also used the chain R of 3.3Å cryo-EM structure of the active, Gs-bound human Calcitonin gene-related peptide (hCGRP) receptor complex (PDB id: 6E3Y) [29] which was the 4th in the hit list. The other two templates we considered included agonist (cholesterol)-bound 3.2 Å structure of human Smoothened receptor (PDB id: 5L7D, ranked 17th in the hit list).

The largest number of hits as potential templates for cAR1 modelling was from class A family of GPCRs. We used four such hits which have been solved in active or active-like conformations. These include: chain A of the activated turkey β_1_ adrenoceptor with a bound agonist (formoterol) and nanobody (PDB id: 6IBL, 2.7Å); chain A of the agonist (NECA-bound) 4.11Å cryo-EM structure of the adenosine A_2A_ receptor complexed with a miniGs heterotrimer (PDB id: 6GDG) [30]; chain R of 3.2Å Crystal structure of the β_2_ adrenergic receptor-Gs protein complex (PDB id: 3SN6) [31] and the chain A of 3Å crystal structure of the thermostabilised human A_2A_ receptor bound to adenosine (PDB id: 2YDO).

It is worthy of mentioning that in DELTA-BLAST searches carried out for cAR2 (uniprot id: P34907), cAR3(uniprot id: P35352) and cAR4 (uniprot id: Q9TX43), members of the class B family of GPCRs ranked similarly high.

### 3.2 Quality of the homology models of cAR1

Using the aforementioned templates, we built seven homology models of cAR1 (arbitrarily named as model A - model G) using Modeller [15]. Although models as stated in the method section were chosen based on the z-DOPE score implemented in Modeller, their subsequent refinement using MD simulation was indeed useful as models after 5ns of MD simulation appeared to be significantly improved in overall 3D quality when evaluated through MolProbity server [17] (Table 1).

**Table 1:** Parameters indicative of the 3D quality of the homology models of cAR1.

| Model name | Template used for homology modelling (PDB id) | 3D quality parameters |  |  |  |  |  |
| --- | --- | --- | --- | --- | --- | --- | --- |
|  |  | Clash score |  | % of poor rotamers |  | % residues in Ramachandran favoured |  |
|  |  | before MD | after MD | before MD | after MD | before MD | after MD |
| A | 6E3Y | 12.2 | 1.7 | 3.1 | 0 | 92 | 96.4 |
| B | 5UZ7 | 10.5 | 1.2 | 1.3 | 0.4 | 94.2 | 98.4 |
| C | 5L7D | 19.8 | 2.7 | 2.2 | 0.4 | 91.7 | 97 |
| D | 3SN6 | 10.8 | 1.9 | 0.9 | 0 | 92.8 | 96.4 |
| E | 6IBL | 17.1 | 1.8 | 0.9 | 0 | 91.8 | 95.1 |
| F | 2YDO | 12.2 | 1.9 | 1.3 | 0 | 93.6 | 96 |
| G | 6GDG | 19.8 | 2.8 | 0.5 | 0 | 92.4 | 97.2 |

### 3.3 Validation of the docking protocol

Prior to predicting the putative cAMP binding site(s) on modelled cAR1 using docking, we first sought out to evaluate the performance of our docking protocol. For this, we chose published structures of 6 different proteins that were elucidated in complex with cAMP. These include the *E.coli* cAMP receptor protein (PDB id: 1I5Z, 1.9Å); the human hyperpolarization-activated cyclic nucleotide-gated ion channel (HCN1, PDB id: 5U6P, 3.51Å); human cyclic nucleotide-binding (CNB) domain of protein kinase A regulatory subunit type Iα (PKA-RIα, PDB id: 5KJX, 1.56Å), the *Mycobacterium tuberculosis* cAMP Receptor Protein, MtbCRP (PDB id: 3I54, 2.2Å); human phosphodiesterase 4D (PDE4D, PDB id: 2PW3, 1.56Å), and mouse exchange protein directly activated by cAMP 2 (Epac2, PDB id: 4MGK, 2.7Å). The bound cAMP was removed from each structure and a different molecule of cAMP obtained from the PubChem was then blindly docked [32] against the aforementioned protein structures AutoDock 4.2. As can be seen in Figure 1, AutoDock 4.2 in blind docking mode could successfully place cAMP to the experimentally-proven binding pockets in all cases and in 4 out of 6 different target protein structures used, the docked pose of cAMP almost overlaid (r.m.s.d ≤1Å). This was impressive since a docked ligand pose at ≤2 Å r.m.s.d heavy-atom distance away from the original (e.g. crystallographic) pose is often regarded as “near-native” solution [33].

**Figure 1.**
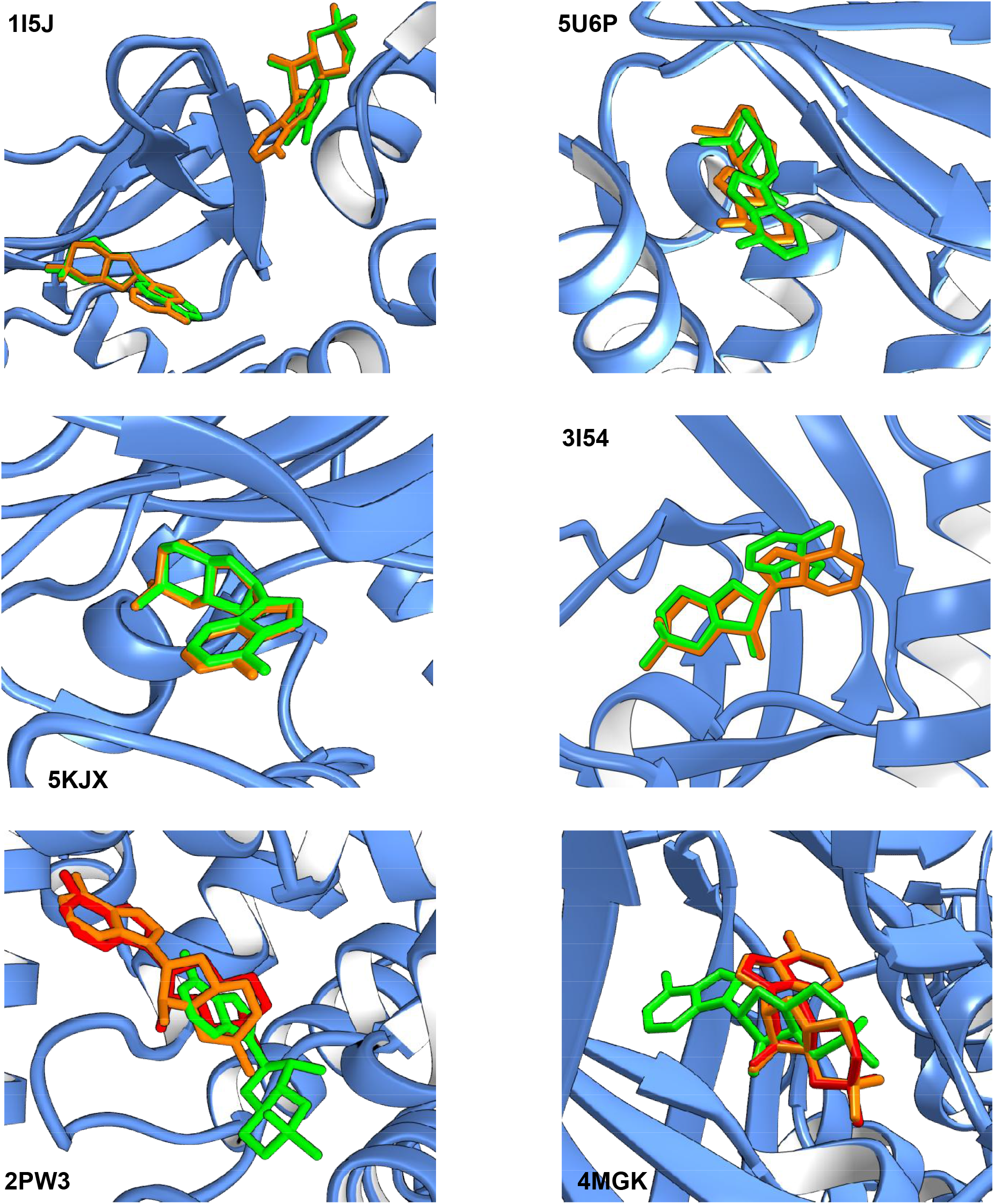
Evaluation of docking protocol in predicting the binding site and binding pose of cAMP on known targets. AutoDock 4.2 was used to blindly dock cAMP on the unliganded structures of several proteins that have been elucidated in complex with cAMP. In set shows the PDB id of each protein shown in a cartoon format whilst the original and docked pose of cAMP are shown as orange and green sticks, respectively. In some cases, a focused docking approach using GOLD suite was undertaken to improve the pose and the resultant docked pose of cAMP is shown in red.

However, in case of human PDE4D (PDB id: 2PW3) and mouse Epac2 (PDB id: 4MGK), AutoDock 4.2 could still place cAMP largely to the experimentally-observed cAMP-binding pockets but the binding poses drifted away considerably (r.m.s.d>2Å) from the original poses. Against the cAMP binding sites of these two protein structures identified by AutoDock 4.2 (and which are the real pockets as well), we re-docked cAMP using GOLD suite (CCDC, Cambridge, UK) and the poses obtained through such GOLD-based focused docking were much improved and nearly superimposed (r.m.s.d <1Å) with the original poses. We therefore decided upon using initial blind docking with AutoDock 4.2 followed by pose refinement using GOLD-based focused-docking as our protocol to model the binding site and binding mode of cAMP on cAR1 homology models.

### 3.4 Prediction of cAMP binding site(s) and choosing the best models for cAR1

Off the total 7 homology models (models A-G) made for cAR1, blind docking of cAMP against 3 of them (models C, D and E) resulted in poses that almost exclusively were located within intracellular regions of the receptor (Figure S2). Structures of active human Smoothened receptor, human β2-adrenergic receptor and turkey β1-adrenergic receptor were used as templates to build these models, respectively. Since an intracellular cAMP binding site for cAR proteins is not plausible, we excluded these models from further analysis and went for inspecting the docked cAMP poses for the remaining four models (models A, B, F, G) of AR1 in all of which the putative binding site of cAMP were in regions similar to the orthosteric sites for GPCRs. These models were built using the structures of active human CGRP receptor (model A), human CTR (model B) and two active human A_2A_ receptors (models F and G).

Previous studies [22–24] employing biochemical and radioligand binding assays with cAMP and its analogs modified at various positions led to the proposition that cAMP binds to the cARs via hydrogen bonds at the amino group (−NH_2_) at C6 position of the adenine ring and at O3’, and that the adenine moiety is likely to be contained within in a hydrophobic cleft of the receptor. We therefore sought out to inspect residue level interaction patterns of the docked cAMP poses for the cAR1 models A, B, F and G using 2D ligand interaction diagrams generated through PoseView™ [27].

In the top-ranked pose of cAMP docked on cAR1 model (model A) based upon CGRP receptor structure as a template (PDB id: 63EY), the adenine moiety seems to project upward whilst the cyclic phosphate group faces downward (Figure 2). Ligand interaction diagram of this pose reveals several hydrogen bonding interactions: between −OH of Ser90 (part of the transmembrane helix 3, TMH3) and the cyclized phosphate group, between −OH of Thr87 (of TMH3) and 3O’; between protonated N1 and −OH of Ser155 (part of the extracellular loop 2, ECL2); between amide carbonyl (>C=O) oxygen of Asn239 (part of TMH7) and the -NH2 at C6 and between Gly153 (part of ECL2) and N7. In addition, hydrophobic interactions are noted between Val154 (part of ECL2) and the diazole sub-ring of the adenine moiety as well as between Phe161 (part of ECL2) and the pyrimidine sub-ring of the adenine moiety. In this way, the docked cAMP seem to make use of the right kind of chemical interactions predicted by previous experimental studies [22–24]. A large difference in affinity for cAMP is known to exist between cARl (25-230 nM) and cAR2 (>5μM). Previous biochemical studies with cAR1 protein mutated at various positions as well as domain-swapped chimera with cAR2 [10, 34] indicated that a part (the second half) of the ECL2 to be mainly accountable for such affinity difference. Among other regions, mutations in TMH3 (Ser90R and Thr97A) also resulted in substantial reduction in cAMP binding affinity [34]. Intriguingly, all the residues that seem to interact with cAMP in the observed pose in model A are part of the ECL2 and TMH3. Thus the observed binding pose of cAMP on model A largely captures the type of interactions predicted by previous biochemical experiments, from both the ligand [22–24] as well as receptor point of view [10, 34]. It is also noteworthy that this cAMP-bound cAR1 model complex was associated with the best docking scores and MM/GBSA energy profile among all other cAMP bound cAR1 model complexes (Table 2).

**Figure 2.**
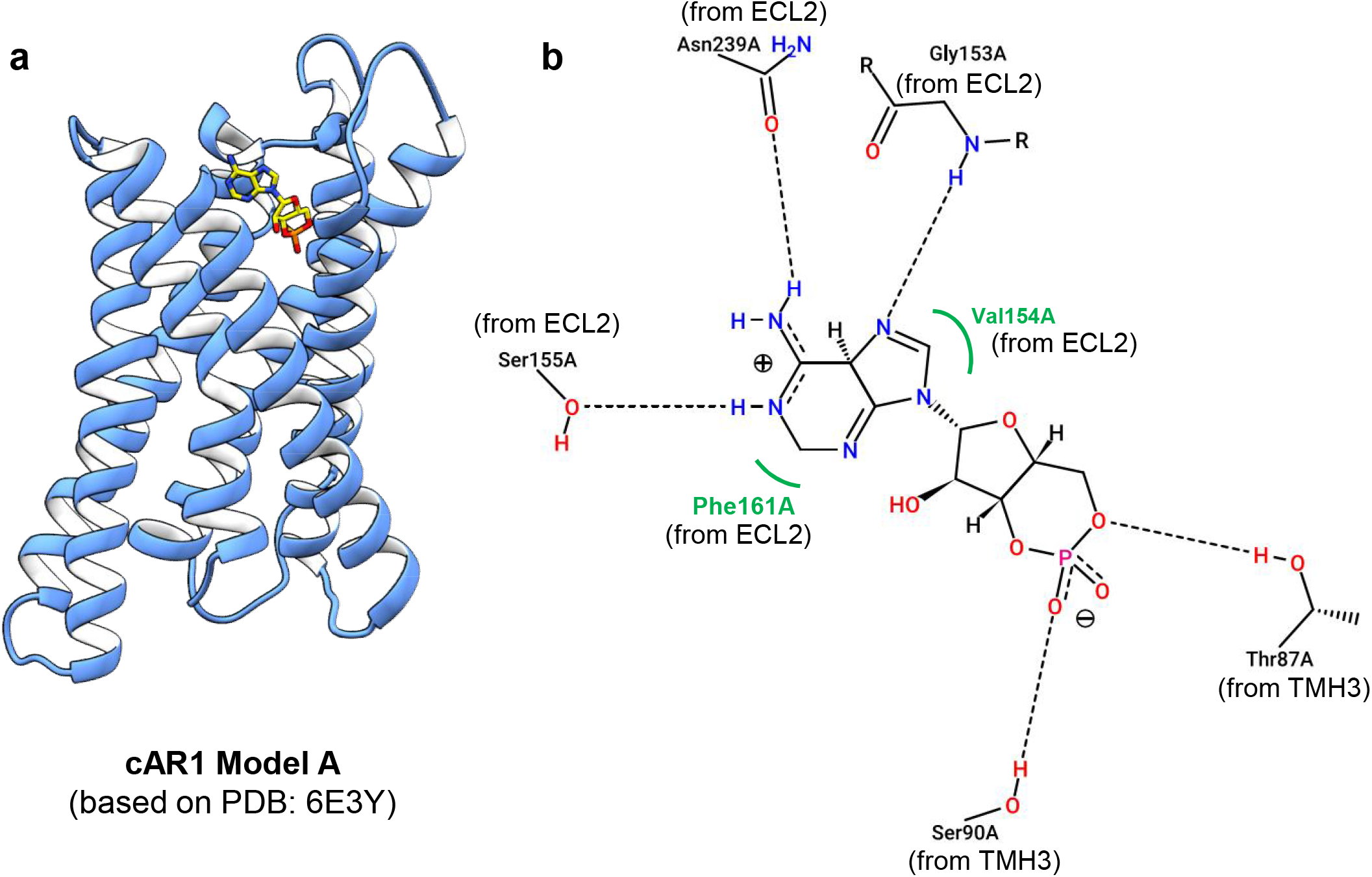
Binding mode of cAMP on cAR1 model A. **A)** The cAR1 model and the docked cAMP are shown as cartoon and stick representations, respectively. Atoms of the cAMP were colour-coded as: blue, N atom; yellow, C atom; red, O atom and orange, P atom. The cAMP pose was produced by initial blind docking using AutoDock 4.2 followed by subsequent pose refinement using GOLD 5.3 suite. **B)** 2D ligand interaction diagram of the cAMP pose shown in left panel. The diagram was generated using Poseview™. Letter A stands for chain ‘A’ designated automatically for the cAR1 model by Poseview. ECL2 and TMH3 stand for extracellular loop 2 and transmembrane helix 3, respectively.

**Table 2:** Parameters related to docking of cAMP of different models of cAR1.

| **cAR1 model used** | **Docking score\*** | | **MM/GBSA rescoring**<br>($\Delta G_{\text{bind}}$, kcal/mol) |
| --- | --- | --- | --- |
| | **AutoDock Vina**<br>($\Delta G$, kcal/mol) | **GOLD**<br>(ChemPLP) | |
| A | -7.8 ± 0.1 | 49.8 ± 0.8 | -47.8 |
| B | -7.8 ± 0.8 | 45.6 ± 0.8 | -46.2 |
| C | -6.9 ± 0.8 | 44.7 ± 2.5 | -38.2 |
| D | -6.2 ± 0.5 | 38.4 ± 1.0 | -30.8 |
| E | -6.4 ± 0.2 | 41.5 ± 1.4 | -34.7 |
| F | -6.4 ± 0.1 | 37.6 ± 1.1 | -21.1 |
| G | -6.1 ± 0.6 | 43.4 ± 1.8 | -24.3 |

\*data represent mean ± s.e.m from 10 independent docking runs

In the top-ranked pose following docking against the cAR1 model (model B) based upon CTR structure as a template (PDB id: 5UZ7), cAMP was placed almost horizontally (relative to the membrane) in a location that was considerably deeper than the orthosteric ligand binding sites observed for GPCRs (Figure 3). In this mode, three hydrogen bonding type interactions are discernible in the corresponding ligand-interaction diagram. One of the phosphate oxygens seems to form H-bonding with the −OH group of Ser246 (of TMH7) whilst the other oxygen seems to be engaged in two different H-bonding interactions with the −OH group of Thr243 (of TMH7) and the guanidinium group of Arg226 (of TMH6). In addition to these, the pyrimidine sub-ring of the adenine moiety seems to be in hydrophobic interactions with Leu66 (of TMH2) and Ile86 (of TMH3) whilst the diazole sub-ring is in hydrophobic contact with Leu17 (of TMH1). This model complex appears to be the second-best model after model A in terms docking score and MM/GBSA energy profiles (Table 2). The pose however, lacks any discernible interaction at the 3O’ position of cAMP as hinted by previous experimental studies [22–24].

**Figure 3.**
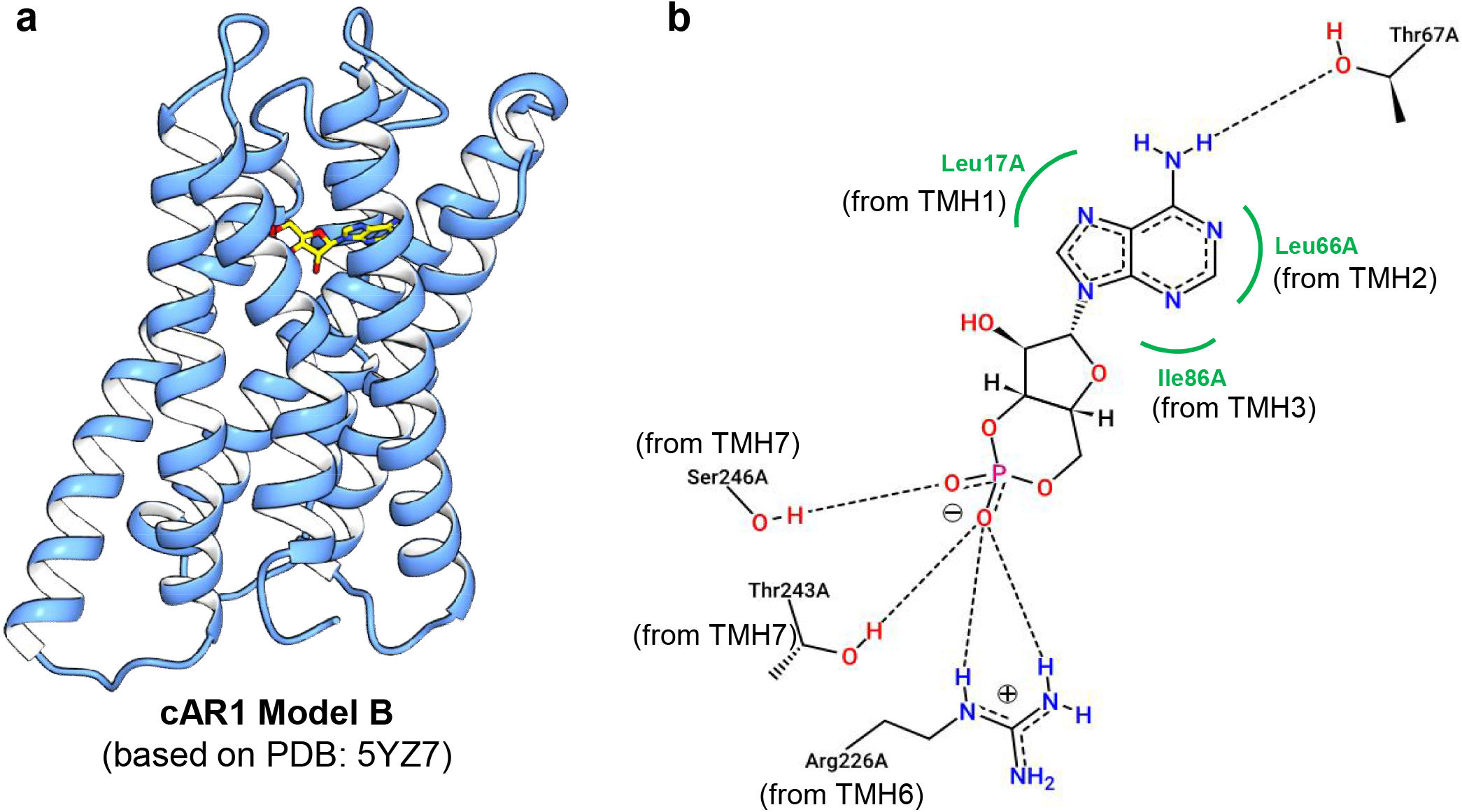
Binding mode of cAMP on cAR1 model B. **A)** The cAR1 model and the docked cAMP are shown as cartoon and stick representations, respectively. Atoms of the cAMP were colour-coded as: blue, N atom; yellow, C atom; red, O atom and orange, P atom. The cAMP pose was produced by initial blind docking using AutoDock 4.2 followed by subsequent pose refinement using GOLD 5.3 suite. **B)** 2D ligand interaction diagram of the cAMP pose shown in left panel. The diagram was generated using Poseview™. Letter A stands for chain ‘A’ designated automatically for the cAR1 model by Poseview. TMH stands for transmembrane helix.

The ligand interaction diagram of cAMP docked on the cAR1 model F (Figure 4) based on A_2A_ receptor (PDB id: 2YDO) shows several H-bonding interactions with residues that almost exclusively belong to the ECL3. For example, there are H bonding interactions between the phosphate group of cAMP and Asn232 (of ECL3), between N7 of the adenine ring and Gly230 (of ECL3) and Leu231(of ECL3) and between the backbone carbonyl oxygen of Arg226 (of TMH6) and the −NH_2_ at C6 of the adenine ring. The pyrimidine subring of the adenine moiety seems to be in hydrophobic interaction with Leu231(of ECL3). Compared to the pose at cAR1 model F, the docked pose of cAMP on the cAR1 model G (Figure 5) which was based on another human A_2A_ receptor structure in complex with miniGs heterotrimer (PDB id: 6GDG) was largely inverted with the adenine moiety facing upward. The cyclized phosphate group is H-bonded with −OH groups of Tyr82 (of TMH3) and Thr243 (TMH7) whilst another H-bonding interaction is observed between 3O’ and Asn239 (of TMH7). However, no H-bonding or hydrophobic interaction was evident with any part of the adenine ring. As a whole, neither models agree with previous experimental suggestions about cAMP binding to cAR1 [10, 22, 23, 34]. The docking scores but the MM/GBSA energies in particular for the docked cAMP poses at these two cAR1 models based on human A_2A_ receptor structures are also substantially lower than models A and B (Table 2).

**Figure 4.**
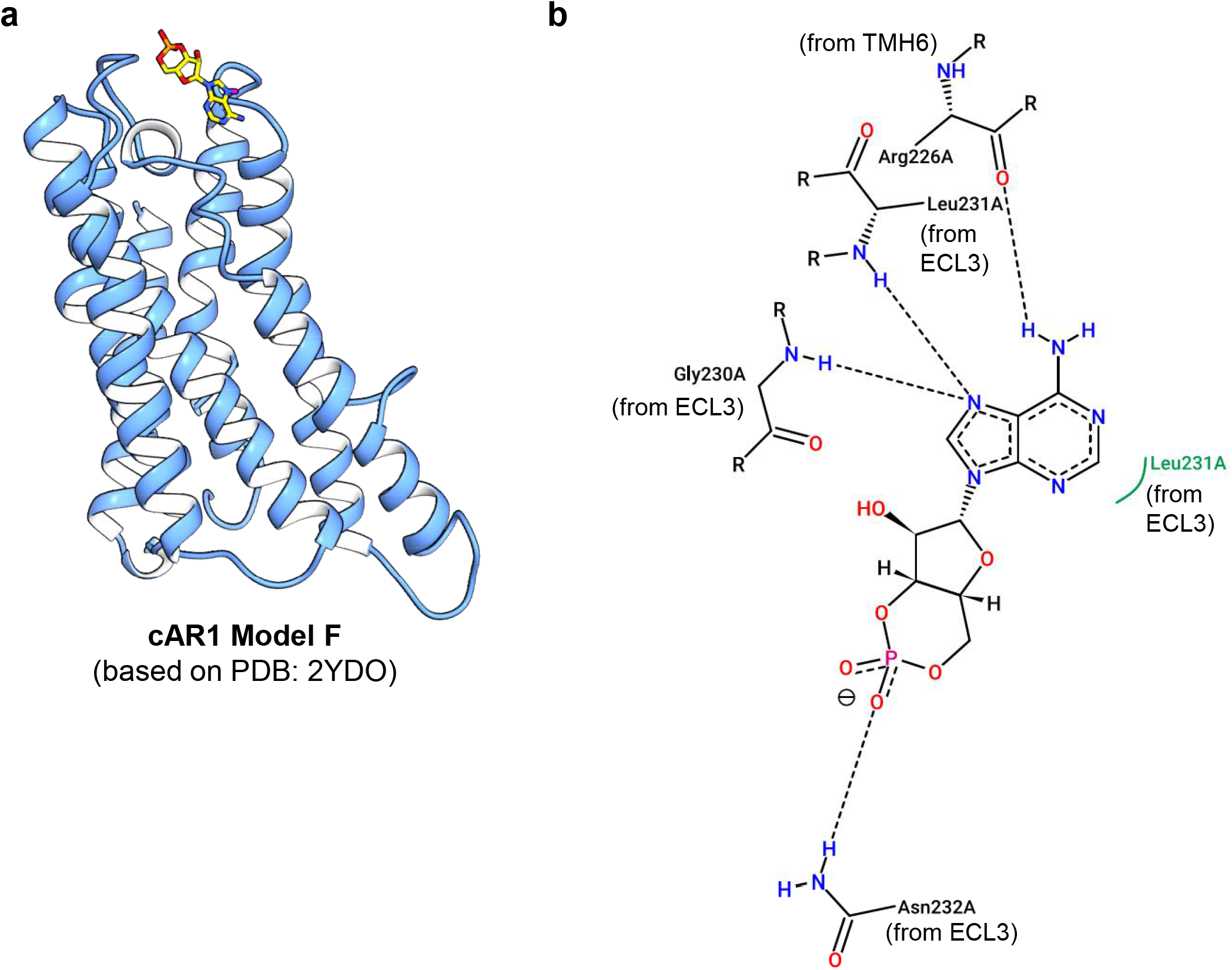
Binding mode of cAMP on cAR1 model F. **A)** The cAR1 model and the docked cAMP are shown as cartoon and stick representations, respectively. Atoms of the cAMP were colour-coded as: blue, N atom; yellow, C atom; red, O atom and orange, P atom. The cAMP pose was produced by initial blind docking using AutoDock 4.2 followed by subsequent pose refinement using GOLD 5.3 suite. **B)** 2D ligand interaction diagram of the cAMP pose shown in left panel. The diagram was generated using Poseview™. Letter A stands for chain ‘A’ designated automatically for the cAR1 model by Poseview. ECL3 and TMH6 stand for extracellular loop 3 and transmembrane helix 6, respectively.

**Figure 5.**
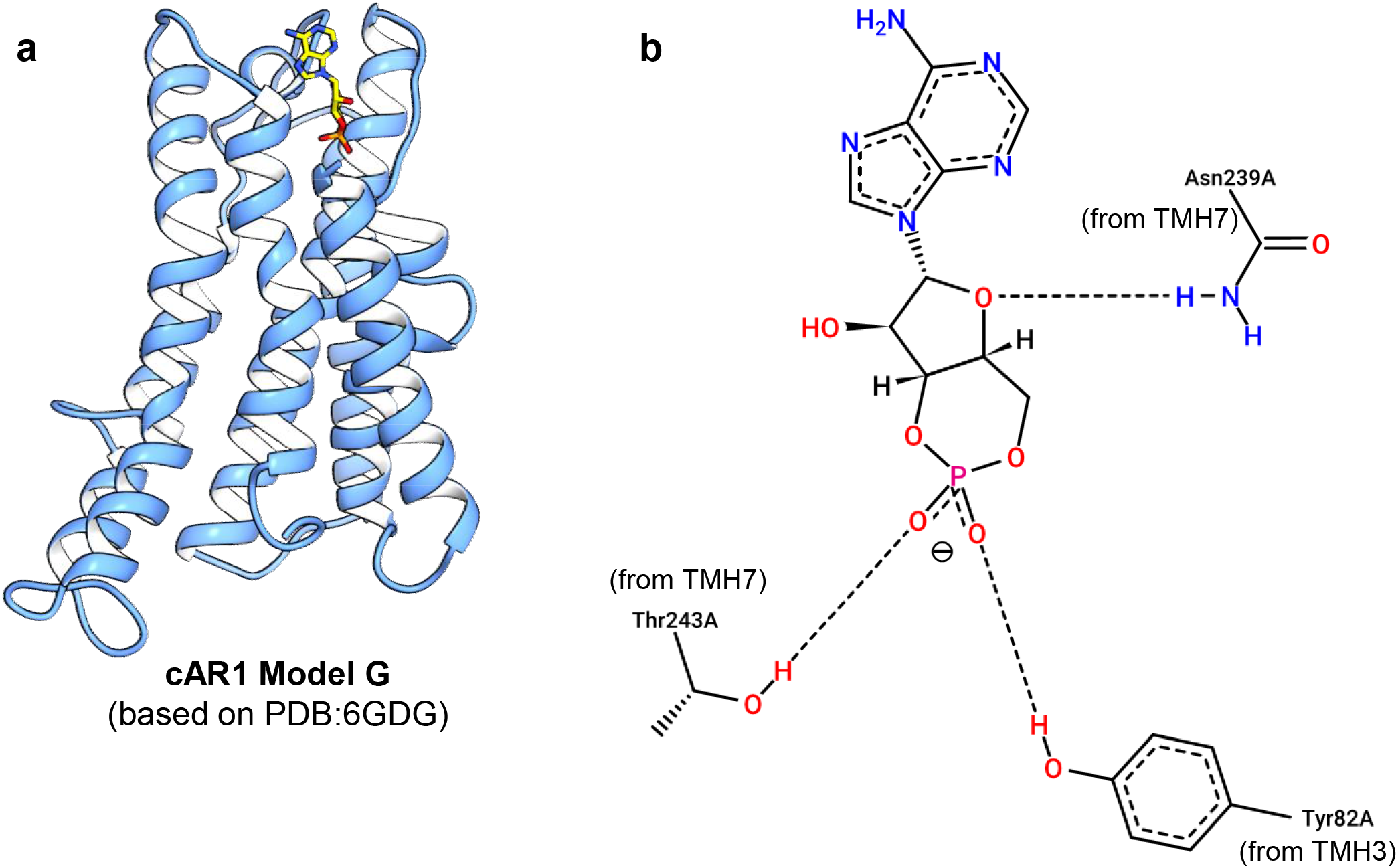
Binding mode of cAMP on cAR1 model G. **A)** The cAR1 model and the docked cAMP are shown as cartoon and stick representations, respectively. Atoms of the cAMP were colour-coded as: blue, N atom; yellow, C atom; red, O atom and orange, P atom. The cAMP pose was produced by initial blind docking using AutoDock 4.2 followed by subsequent pose refinement using GOLD 5.3 suite. **B)** 2D ligand interaction diagram of the cAMP pose shown in left panel. The diagram was generated using Poseview™. Letter A stands for chain ‘A’ designated automatically for the cAR1 model by Poseview. TMH stands for transmembrane helix.

With a view to finding further aid for selecting the most plausible cAR1 model(s), we sought out to predict the binding pose and energy of two analogs of cAMP namely 3’-Deoxy-3’-aminoadenosine 3’, 5’-monophosphate (3’-NH-cAMP) and 6-Chloropurineriboside 3’,5’-monophosphate (6-Cl-PuRMP) (Figure S1) against all the cAR1 models A, B, F and G. Both these compounds have been experimentally proven to show much reduced affinity to bind cAR1 presumably due to lack of any H-bonding interactions at the 3’O and 6-NH_2_ positions [22, 24]. As can be seen in supplementary table S1 (Table1 S1), both 3’-NH-cAMP and 6-Cl-PuRMP docked to model A roughly to the same pocket but with significantly reduced scores with AutoDock 4.2 and ChemPLP score (GOLD) as well as reduced binding energy estimated by MM/GBSA. Close inspection of the poses revealed that indeed no H-bonding interaction with any residue could take place at the modified positions (3O’ and C6) and the some changes in the binding pose specially for the cyclized phosphate moieties were also noted with respect to the cAMP pose at this pocket for model A (Figure 6). Thus, H-bonding at these two positions of cAMP seems to be a major contributing factor governing ligand binding affinity for the cARs and this largely agrees with previous experimental suggestions [22–24].

**Figure 6.**
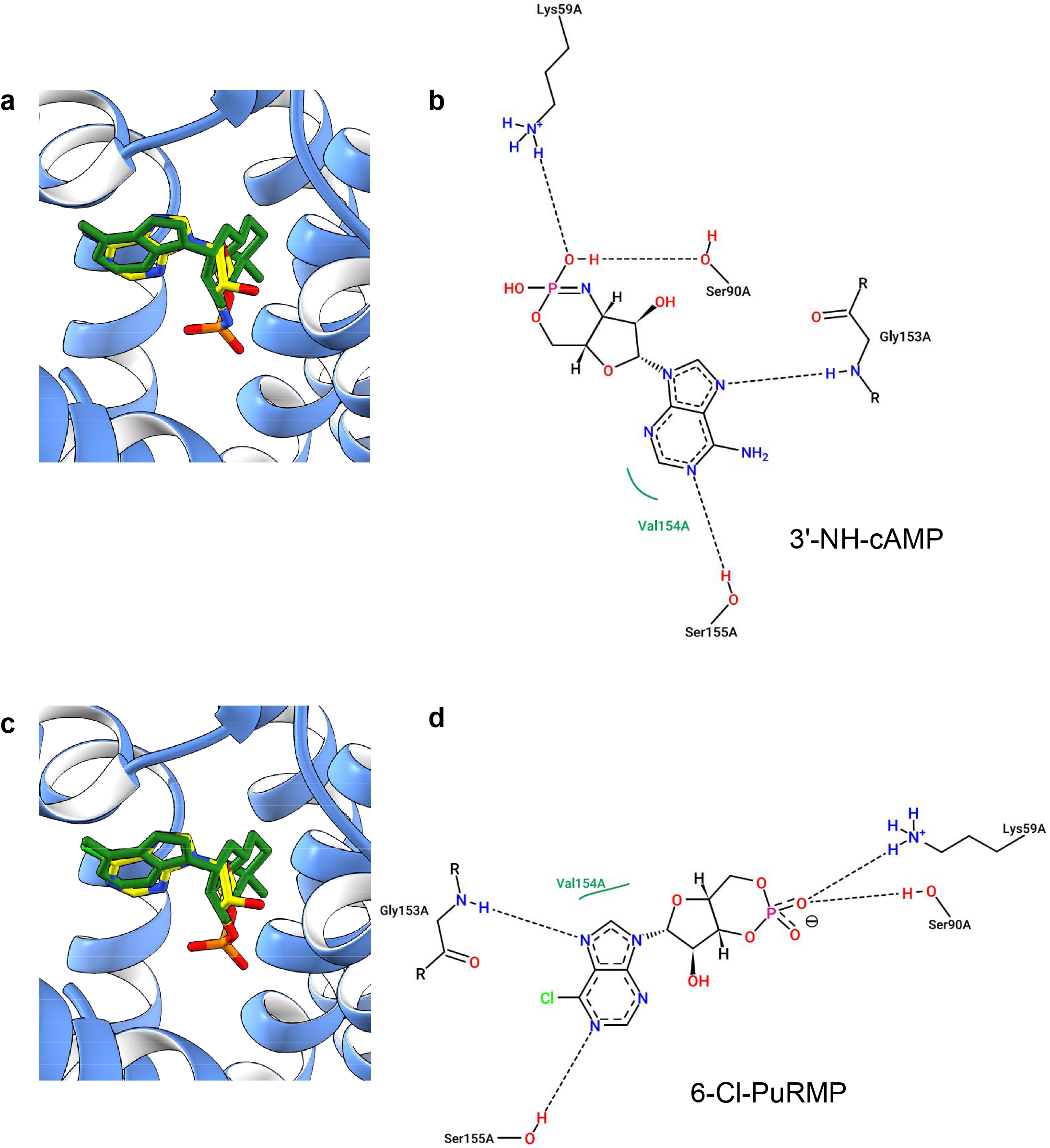
Binding mode of cAMP analogs on cAR1 model A. Left panels **A** and **C** show model A with docked 3’-NH-cAMP and 6-Cl-PuRMP, respectively. The cAR1 model and the docked cAMP analogs are shown as cartoon and stick representations, respectively. Atoms of the cAMP analogs were colour-coded as: blue, N atom; yellow, C atom; red, O atom; orange, P atom and light green, Cl atom. Docked cAMP pose are shown as deep green sticks. All ligand poses were produced by initial blind docking using AutoDock 4.2 followed by subsequent pose refinement using GOLD 5.3 suite. The right panels **B** and **D** show the 2D ligand interaction diagrams for the docked complexes shown in left panels. The diagram was generated using Poseview™.

For model B, the consequence of modifying cAMP at the 3’ and 6 position appeared to be more drastic since both 3’-NH-cAMP and 6-Cl-PuRMP preferentially docked to intracellular faces of the receptor. It was therefore not feasible to compare the corresponding docking score to compare with the cAMP pose (data not shown). For models F and G which were built based on two different active structures of human A_2A_ receptor, no substantial change in binding energy (Table S1) or poses (not shown) were noticed.

Based on type of interactions of cAMP at the residue levels, the docking scores and MM/GBSA energy profiles and the consequences observed for the cAMP analogues modified at two key positions on binding pose and score/energy, model A definitely stands out as the most plausible of all and model B is perhaps the second best in this regard. For both model A and B, we wanted to evaluate the extent of the evolutionary conservation of amino acids, particularly at the putative cAMP binding sites. For this purpose, we used Consurf which uses an empirical Bayesian method to compute position-specific conservation scores for each amino acid in the query protein sequence based on the phylogenetic relations between homologous sequences [26]. Among the residues interacting with docked cAMP in model A (Figure 2), Phe161 and Ser90 appear to be strongly conserved; conservation at Thr87, Asn239 and Gly153 was intermediate whilst Val154 and Ser155 appear to be variable (Figure 7A-B) In contrast with model A, the residues interacting with cAMP in model B appear to be well conserved (Figure 7C-D). To compare, we looked into the Consurf-DB (http://bental.tau.ac.il/new_ConSurfDB/) [35] and visually inspected the degrees of evolutionary conservation of residues at the cAMP binding pockets (the CNB domains) of some known cAMP binders including *E.coli* cAMP receptor protein (PDB: 1I5Z), hyperpolarisation-activated cAMP-gated ion channels isoform 2 and isoform 4 (HCN2, HCN4; PDBs: 1QPO and 3OTF, respectively), human phosphodiesterase isoform 4D (PDE4, PDB: 2PW3) and the RI regulatory subunit of PKA (PDBs: 1RGS, 3PNA). Although in general most of the cAMP-interacting residues for these proteins appear to be conserved, the degrees of conservation vary to some extent across protein classes (data not shown). However, it is noteworthy that the residues interacting with the phosphate group tend to be very well conserved whilst those interacting with the adenine moiety manifest some variation in conservation. From this point of view, the evolutionary conservation profiles of cAMP interacting residues in model A deem reasonable.

**Figure 7.**
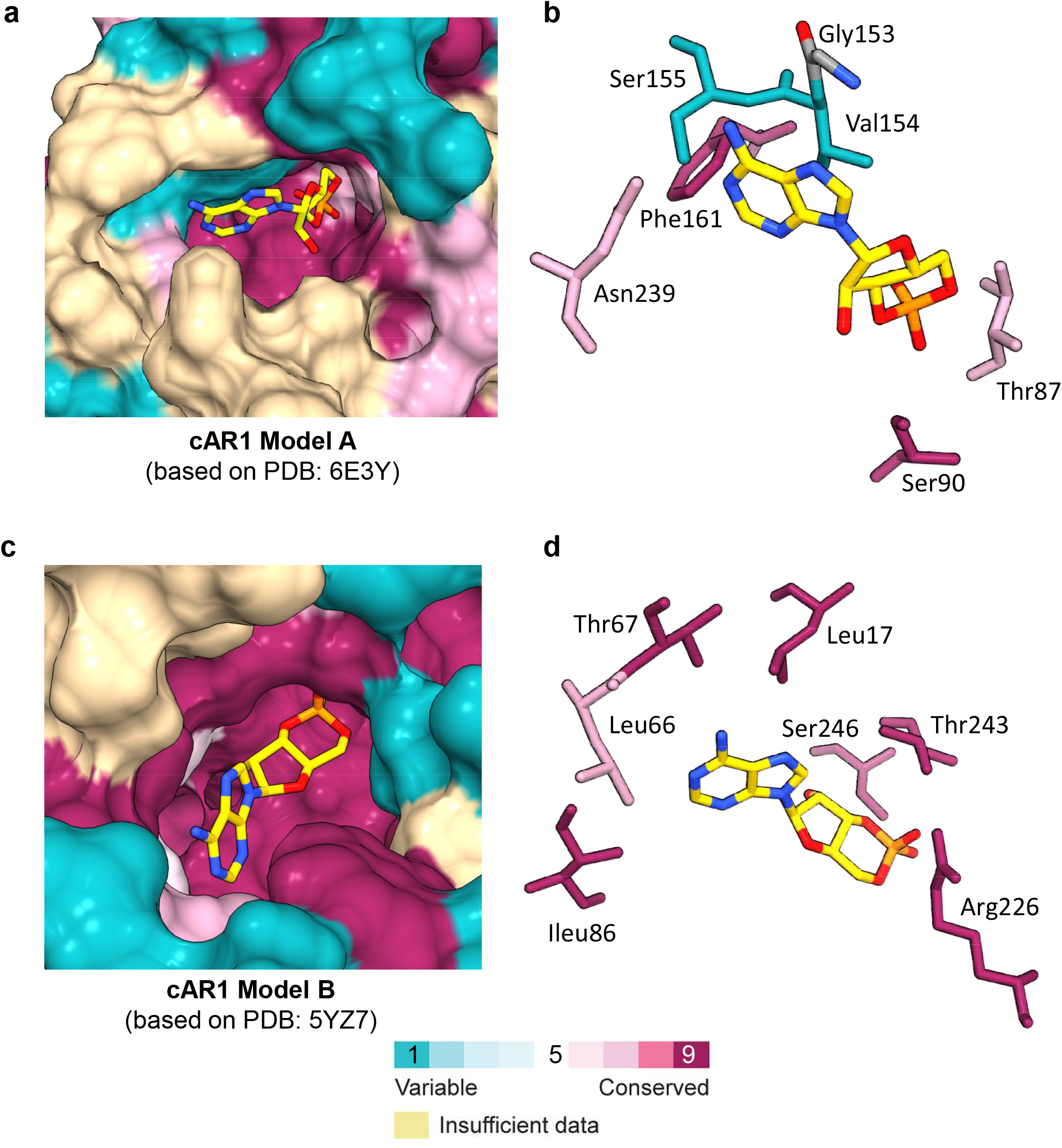
Evolutionary conservation of residues within the putative cAMP binding pockets of models A and B. The left panels, **A** and **C** show the putative binding pockets of cAMP in models A and F shown in surface representations, respectively. In the right panels **B** and **D**, cAMP and the interacting residues relevant to the panels A and B are shown in stick representations. The amino acids of cAR1 models are coloured in the range from turquoise to maroon based on conservation grades according to ConSurf description given below.

### 3.5 Modelling other three cAR proteins bound to cAMP

Since model A of the cAR1 bound to cAMP appears to be a plausible one, we went for using the same parent template (CGRP receptor structure, PDB id: 63EY) and essentially the same homology modelling protocol to model the remaining members of the cAR family namely cAR2, cAR3 and cAR4. We also felt this follow-up modelling task logical since these cAR subtypes manifest significant difference with respect to cAR1 in their affinity for cAMP, in addition to the difference in the spatio-temporal aspects of their expression [5, 6]. All final models of these three cAR subtypes were comparable with cAR1 model A in overall 3D quality (Table S2). Docking of cAMP to these models using the same methodology (blind docking by AutoDock 4.2 followed by focused docking using GOLD and estimation of MM/GBSA energies) revealed broadly the similar location of cAMP binding, however significant difference in some docking scores (Table S3) and poses (Figure 8) were noted.

**Figure 8.**
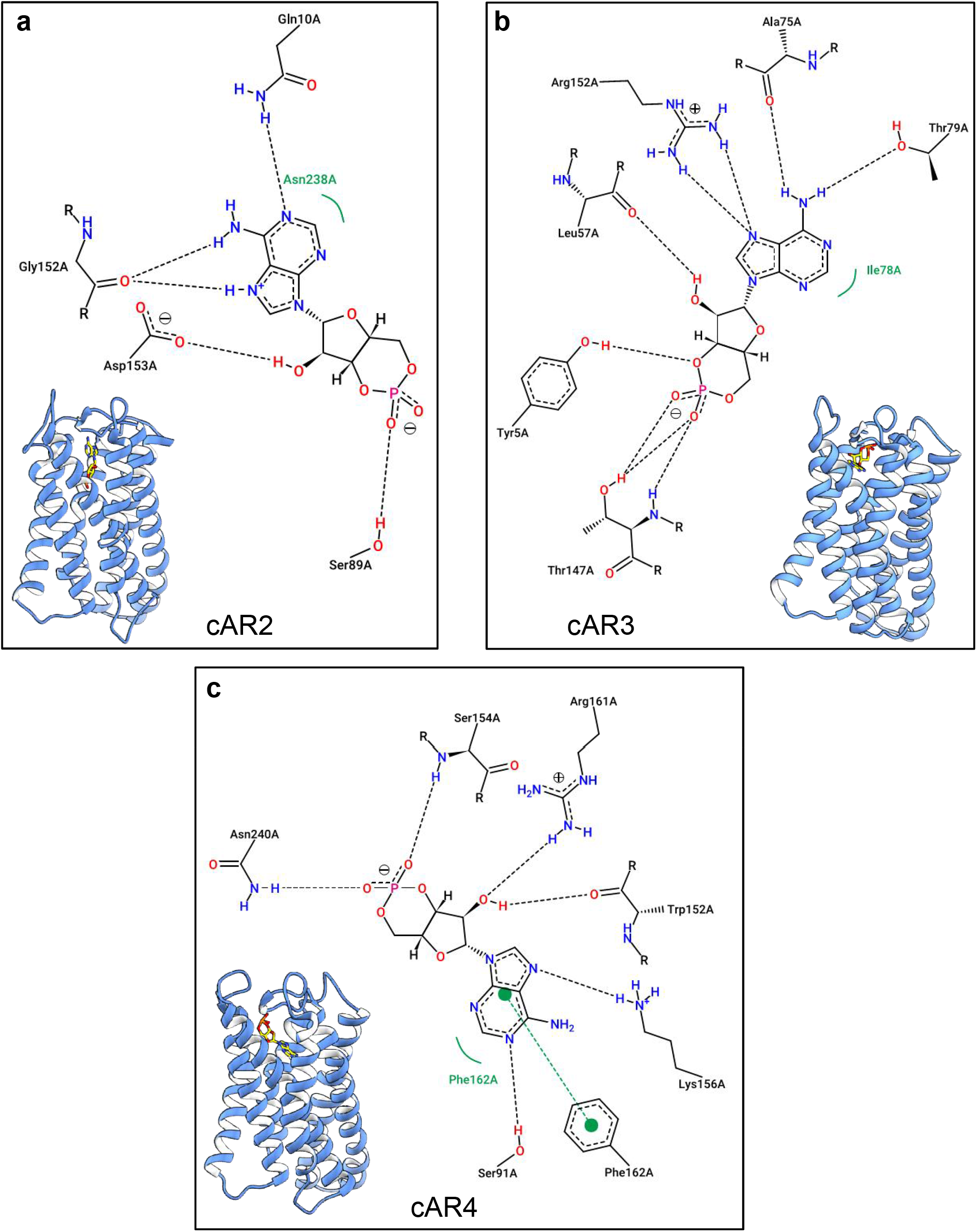
Binding mode of cAMP on models of cAR2, cAR3 and cAR4. **A)** cAR2 model with docked cAMP and the corresponding 2D ligand interaction diagram is shown **B)** as for **A** except cAR3 model **C)** as for **A** except cAR4 model was used. In all cases, cARs are shown as cartoons whilst cAMP poses are represented as sticks with atoms colour-coded as: blue, N atom; yellow, C atom; red, O atom and orange, P atom. The cAMP pose was produced by initial blind docking using AutoDock 4.2 followed by subsequent pose refinement using GOLD 5.3 suite. 2D ligand interaction diagrams were generated using Poseview™.

For cAR2 model, the docked pose of cAMP (Figure 8A) was almost similar to that of cAR1 model A but the cognate docking scores (Table S3) in AutoDock 4.2 and GOLD as well as the MM/GBSA energy were significantly lower than those of the cAR1 model A. Upon close inspection of the corresponding 2D ligand interaction diagram in comparison with of cAR1 model A, the apparent reduction in the docking score and MM/GBSA energy for cAMP pose on cAR2 model seem mainly to stem from loss of H-bonding interaction at 3O’ position with an additional contribution from lack of hydrophobic interaction at the diazole sub-ring of the adenine moiety (Figure 8A). This is intriguing because two cAMP analogues namely 3’-NH-cAMP and 6-Cl-PuRMP that are known to manifest much lower affinity for the cAR1 receptor also seem to miss H-bonding at 3O’ position in addition to position C6 (Figure 6).

The docking scores and MM/GBSA energy profiles for cAMP pose at the cAR3 model were significantly higher than those for cAR2 and cAR4 models (Table S3). It was however not readily clear why this was the case, especially when compared to the cAR2 except the fact that there was a H-bonding occurring 5O’ and Tyr5 which could perhaps compensate for the lack of H-bonding at 3O’ (Figure 8B).

The docking scores and MM/GBSA energy profiles for cAMP pose at the cAR4 model were also low, comparable to those of cAR2 (Table S3) and this could be due to lack of any H-bonding interactions at the 3O’ and C6 position (Figure 8C). It is noteworthy to see a π-π stacking between the adenosine moiety and Phe162.

As mentioned before, previous experimental studies have well established that cAR proteins differ markedly in their binding affinity (K_d_) for cAMP with cAR1 and cAR3 having the highest (~300nM) and second highest (~500nM) for cAMP, respectively the cAMP affinities of cAR2 and cAR3 are substantially lower (>5μM) [5]. Our modelling and docking approach could not exactly recapitulate such differential affinity profile of cARs but we did not expect to be successful in this aspect either. Reliable estimation of binding affinities or free energies using in silico approaches is not trivial, often requiring extensive MD simulation and quantum mechanical calculations and methods are still evolving. In the present study, we did not pursue this aspect thoroughly but relied upon docking scores and MM/GBSA method to predict cAMP poses with a view to selecting the best models of cAR proteins. However, the overall trend that we found, especially based upon the GOLD ChemPLP score and MM/GBSA-derived energies reasonably agrees with the experimentally-observed cAMP-based affinity ranking of cARs [5].

### 3.6 Comparison of the predicted cAMP binding pockets of cARs with the CNB domains

The canonical CNB domain characteristically contains a conserved, eight stranded β barrel domain (β subdomain) within which there is a phosphate binding cassette (PBC) that anchors the phosphate group of cAMP via a conserved Arg (e.g. Arg209 for PKA). This Arg appears to be properly poised to do so through an interaction with a conserved Gly (Gly169 for PKA) present within the β2-β3 loop. The CNB domains also contain an α-helical subdomain that is more variable in sequence and structure [12]. The cARs lack any sequence compatible with the CNB or similar domain, however some analogous features can still be discerned from the 2D ligand interaction diagrams of the cAR-cAMP complexes (Figures 2,8). For example, the phosphate group seems to interact with a Ser and/or Thr in case with the cARs and intriguingly a Ser or Thr is present adjacent to the critical Arg in the PBC of the CNB domain for main proteins [12] and often this Ser/Thr also seem to participate in H-bonding interaction with the phosphate group for some cAMP binding proteins including CAP of *E.coli* (pdb: 4HZF), human HCN4 channel (pdb: 4HBN) and human PKG (pdb: 5C6C). The adenine moiety of cAMP is accommodated largely by some hydrophobic residues within the β domain (mainly β6 and β7 sheets) of the CNB whilst the amino group at C6 of cAMP form H-bonding with some residues in some cases (for e.g. human Epac1/2,pdb: 4MGI or 4MGK; *E.coli* CAP, pdb: 4HZF). Broadly similar types of interactions can be discerned for the adenine moiety of cAMP docked onto our cAR1, cAR2 and cAR3 models (Figures 2,8). For our cAR4 model, although a Ser participates in H-bonding with the phosphate group, there is no such interaction occurring for the amino group at the C6 position and the adenine moiety seems to make a π-π stacking interaction with a Phe (Figure 8C). Interestingly, such π-π stacking with the adenine ring is observed for the human A_2A_ receptor structures bound to adenosine (pdb: 2YDO) and NECA (pdb: 5G53, 6GDG, 2YDV) as well as for few proteins with the CNB domain that notably include PDEs (pdb: 2PW3, 2OUR) and regulatory subunit of PKA (pdb: 1RGS). Thus, overall analyses based on the 2D ligand interaction diagrams of docked cAMP poses on our cAR models reveal plausible structural arrangements for cAMP recognition, even though there is clearly no canonical CNB or similar domain present in these GPCRs.

### 3.7 Structural comparisons between the cARs and other GPCRs

It is also worthy of mentioning that none of the conserved microswitches (e.g. the E/DRY motif in TMH3, the CWxP motif in TMH6, the NPxxY motif at the intracellular end of TMH7) that are well known to play important role in mediating the activation mechanism of many GPCRs [36] were clearly evident in the sequences or in our homology models of cARs. These microswitches are mostly established for class A family of GPCRs whilst they appear to be absent in class B GPCR family members including CGRP receptor which served as the best template for modelling cARs. In some cases, there can be positional equivalents of some of the abovementioned microswitches and few other class A GPCR motifs detectable in class B GPCR family members, but their functional significance may not be comparable to those for the class A GPCRs [37, 38]. For example a triad of conserved residues in TMH3 (^236^YLH^238^ for hCGRP receptor, pdb: 6E3Y) of class B1 family of GPCRs has been suggested to be a positional but not functional equivalent of the E/DRY motif present in most members of the class A family [37]. However, mutations in this motif for class B seem not to affect receptor function as markedly as seen for class A members mutated at the E/DRY motif although a double Ala mutant (^236^YL →AA) had ~70% less surface expression [39]. In a comparable position, cARs seem to have a conserved Ser-Ile-Try (^103^SIY^105^ for cAR1) (Figure S3).

At a location just upstream of ICL2, a pair of aromatic amino acids (^254^WY^255^ for hCGRP receptor, pdb: 63EY) appear to be noticeably conserved within TMH4 across most of the class B family members and these residues can be structurally and functionally important [38]. For example, mutation at Trp254 (^254^W) has been shown to reduce CGRP potency for hCGRP receptor whilst Tyr255 (^255^Y) is among several residues that interact the Receptor activity-modifying protein 1 (RAMP1) with a possibility of forming H bonding with S141 of RAMP1 [29]. Interestingly, we notice comparable aromatic residues (Tyr/Phe, Tyr) at equivalent position in all our cAR models (e.g. ^121^YY^122^ for cAR2 model A; Figure S3, Figure S4A).

Another tetrad of residues, namely the KKLH (or comparable) motif in ICL1, is shared between class A and class B family [37]. For the hCGRP receptor structure (pdb: 6E3Y) which served as the template for our finally-chosen models of cARs, the sequence in the analogous position is ^167^KSLS^170^. Intriguingly in our models of cARs, we could find comparable sequence ‘KLLR’ in analogous location: ^39^KLLR^42^ for cAR1 model A, ^36^KLLR^39^ for cAR2 model, ^30^KLLR^32^ for cAR3 model and ^36^KLLR^39^ for cAR4 model (Figure S3, figure S4B). Upstream of ICL1, an Arg (R173 for hCGRP receptor structure, pdb: 6E3Y) located at the bottom end of TMH2 appears to be well conserved across all family B GPCRs [38] and an Ala mutant at this position for CGRP receptor is known to cause severe disruption in associated signalling towards cAMP production [39]. However, instead of an Arg or (any polar residue), a Val/ Ile flanked by conserved His and Thr can be seen across cARs sequences (Figure S3) but their functional significance remains unknown.

Although the NPxxY^7.53^ motif (or its equivalent) typically observed in class A GPCRs is absent in cARs, a conserved, positional equivalent of the Y^7.53^ could be located in our cARs models: F257 for cAR1 model A, F256 for cAR2, F249 for cAR3 and F258 for cAR4 model (Figure S4C).

Last but not the least, a highly conserved GWGxP motif (^259^GWGFP^263^ for hCGRP receptor, pdb: 63EY) within the TMH4 is present in all class B GPCRs [38] and this motif is known to play important structural role in stabilizing the configuration of TMH2, TMH3 and TMH4 [40]. Although exact version of such motif is not present in cARs, a potential positional equivalent is noticeable with a Cys replacing the initial Gly (^126^CWGLP^130^ for hCGRP receptor structure, pdb: 6E3Y).

Thus there appears to be some comparable structural features between the cARs and the class B family of GPCRs and this may help in explaining why a class B GPCR served as the best template for modelling these class E family members. However, the functional significance of all the aforementioned residues of cARs need to be experimentally validated in future studies.

## Conclusion

Using ligand docking-guided homology modelling approach, we have constructed plausible models of the active form of all four cAR proteins of *D. discoideum* - a widely used organism for studying chemotaxis and developmental signalling. Our analysis indicates an orthosteric cAMP binding domain that is structurally distinct from the canonical CNB domains but retains broadly similar type of interactions at residue levels. Although it remains technically still challenging to reliably predict ligand-binding affinities through in silico approaches, our models nevertheless could successfully recapitulate the overall trend of differential cAMP binding affinities that are well established across the cAR subtypes. We believe our models could prompt experimental validation studies of cAMP recognition at cAR proteins, involving site directed mutagenesis, radio-ligand binding and functional assays in future.

## Supporting information

supplementary_blast_search

---

**Figure S1. 2D chemical representation of cAMP and two analogues used for docking against cAR1 models**

**Figure S2. Binding mode of cAMP on selected models of cAR1**. Blind docking using AutoDock Vina was primarily used to identify the most likely binding site of cAMP whilst the binding mode was subsequently refined through focused docking using GOLD 5.3 suite.

**Figure S3. Sequence alignment of the modelled regions of cAR isoforms.**

The primary sequences of cAR1 (uniprot id: P13773, residues: 1-268), cAR2 (uniprot id: P34907, residues: 1-267), cAR3 (uniprot id: P35352, residues: 1-260), P34907 and cAR4 (uniprot id: Q9TX43, residues: 1-269) were aligned using Clustal Omega (https://www.ebi.ac.uk/Tools/msa/clustalo/) in its default setting. Important residues known for class B are shown in red whilst the positional equivalents of those for cARs are shown in blue.

**Figure S4. cAR residues that are potential positional equivalents of known GPCR motifs important for receptor activation.** cAR1 model A (shown in light blue) was superimposed on human CGRP receptor (shown in yellow, pdb: 6E3Y) using UCSF Chimera. Comparable residues are shown as sticks. Panel A shows motif may be positional equivalent of the E/DRY microswitch. Panel B shows the positional equivalent of KKLH (or similar) motif shared between class A and B family of GPCRs. Panel C shows the positional equivalent of Y^7.53^ of the NPxxY^7.53^ motif.

**Table S1:** Parameters related to docking of cAMP analogues against selected models of cAR1.

| ligand | cAR1 model used | Docking score* | | MM/GBSA rescoring<br>($\Delta G_{\text{bind}}$, kcal/mol) |
| --- | --- | --- | --- | --- |
| | | AutoDock Vina<br>($\Delta G$, kcal/mol) | GOLD<br>(ChemPLP) | |
| 3'-NH-cAMP | A | $-6.5 \pm 0.08$ | $31.3 \pm 1.2$ | -30.3 |
|  | B | ND | ND | ND |
| | F | $-6.3 \pm 0.5$ | $45.74 \pm 2.5$ | -23.9 |
| | G | $-6.1 \pm 0.5$ | $38.41 \pm 1.0$ | -25.1 |
| 6-Cl-PuRMP | A | $-6.5 \pm 0.2$ | $40.05 \pm 1.5$ | -31.7 |
|  | B | ND | ND | ND |
| | F | $-6.2 \pm 0.5$ | $43.45 \pm 1.8$ | -23.0 |
| | G | $-6.0 \pm 0.1$ | $49.05 \pm 0.04$ | -26.1 |

\*data represent mean ± s.e.m from 10 independent docking runs; ND, not determined

**Table S2:** Parameters indicative of the 3D quality of the homology models of cAR1, cAR2 and cAR3.

| Model name | 3D quality parameters |  |  |  |  |  |
| --- | --- | --- | --- | --- | --- | --- |
|  | Clash score |  | % of poor rotamers |  | % residues in Ramachandran favoured |  |
|  | before MD | after MD | before MD | after MD | before MD | after MD |
| cAR2 | 5.4 | 2.3 | 0.4 | 0 | 88.3 | 95.5 |
| cAR3 | 3.6 | 1.835 | 1.3 | 0 | 88.1 | 96 |
| cAR4 | 5.4 | 2.3 | 1.8 | 0 | 87.1 | 96.4 |

**Table S3:** Parameters related to docking of cAMP against homology models of cAR2, cAR3 and cAR4.

| cAR model used | Docking score* | | MM/GBSA rescoring<br>($\Delta G_{\text{bind}}$, kcal/mol) |
| --- | --- | --- | --- |
| | AutoDock 4.2<br>($\Delta G$, kcal/mol) | GOLD<br>(ChemPLP) | |
| cAR2 | $-6.2 \pm 0.05$ | $33.4 \pm 1.0$ | -30.57 |
| cAR3 | $-7.0 \pm 0.05$ | $42.4 \pm 1.0$ | -40.13 |
| cAR4 | $-6.5 \pm 0.05$ | $35.4 \pm 1.5$ | -32.4 |

**Figure S1.**
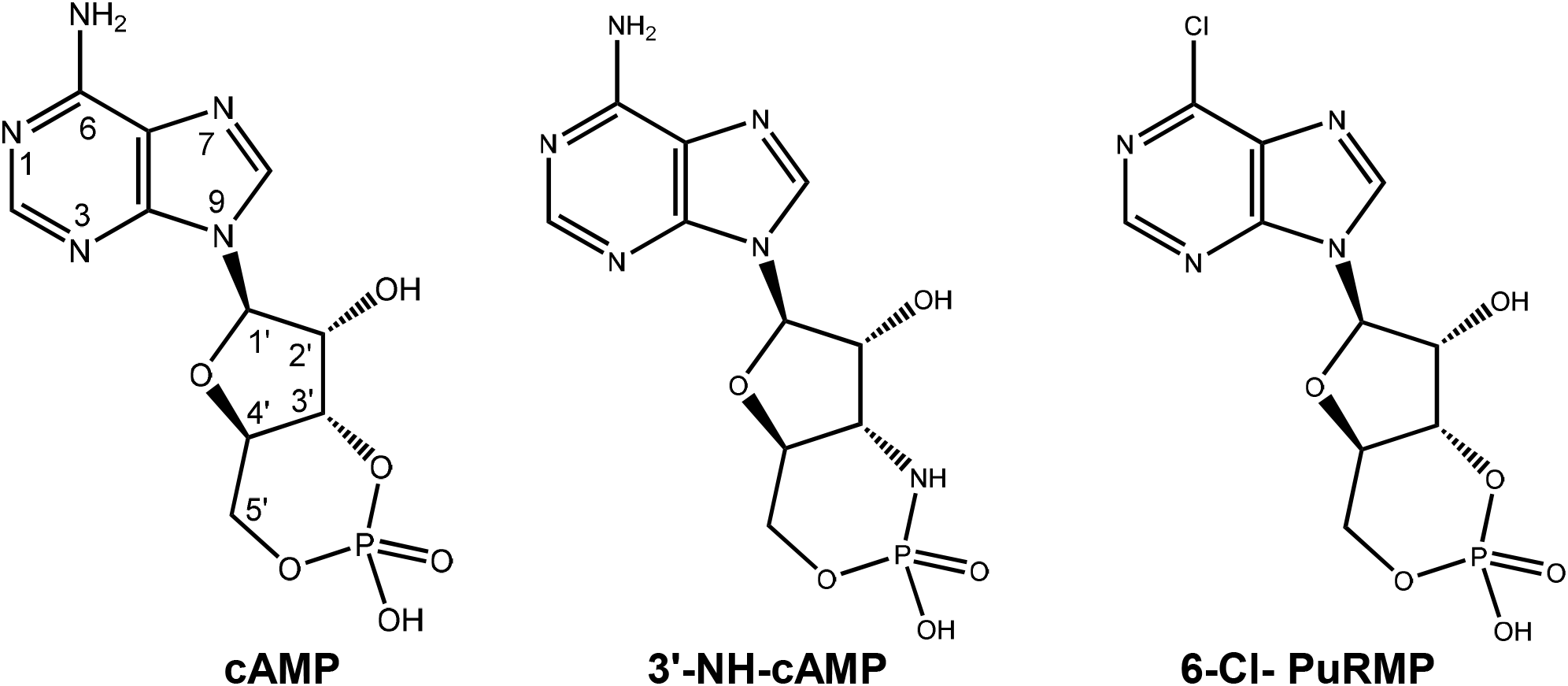
2D chemical representation of cAMP and two analogues used for docking against cAR models in the present study

**Figure S2.**
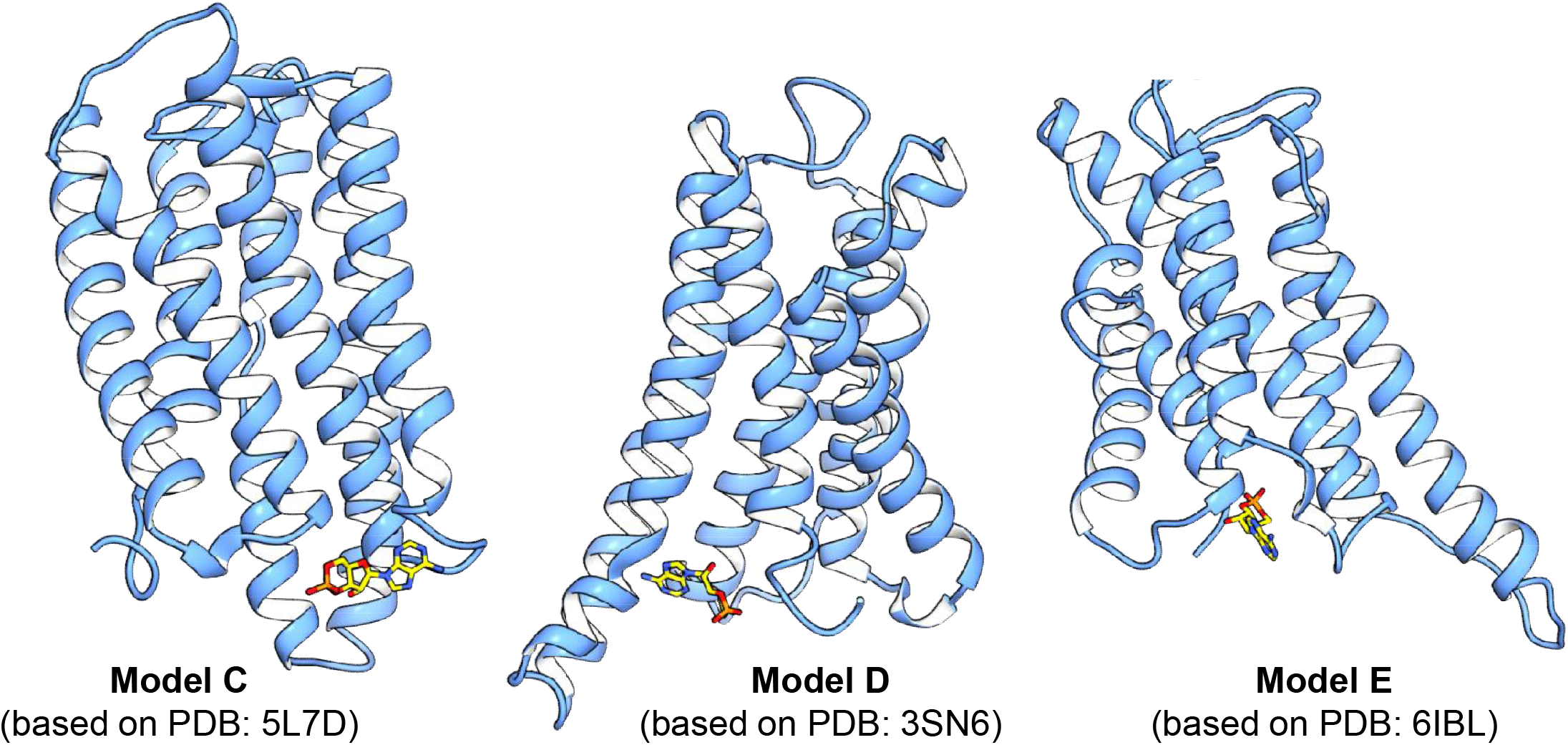
Binding mode of cAMP on selected models of cAR1. Blind docking using AutoDock 4.2 was primarily used to identify the most likely binding site of cAMP whilst the binding mode was subsequently refined through focused docking using GOLD 5.3 suite.

**Figure S3.**
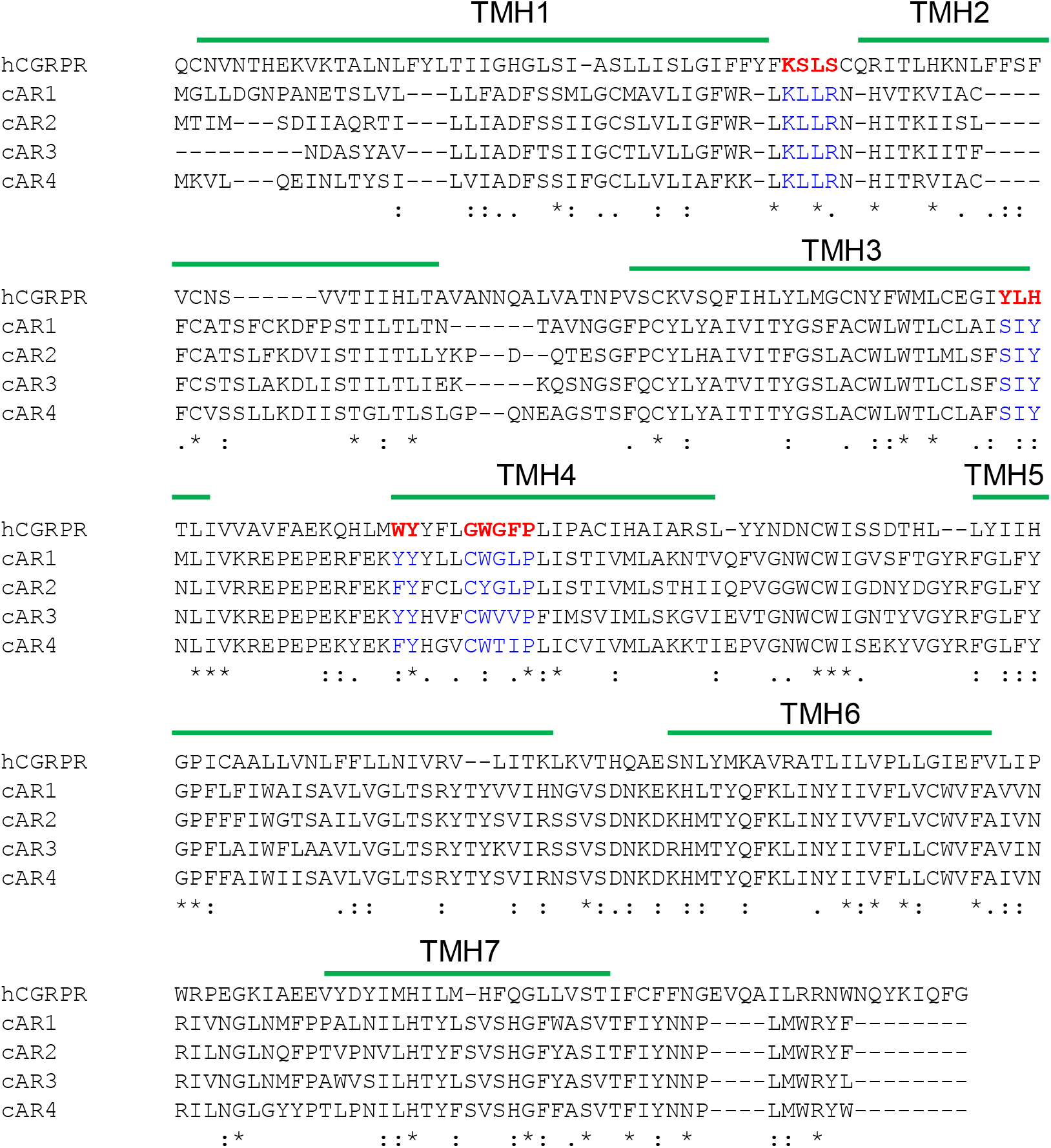
Sequence alignment of the modelled regions of cAR isoforms. The primary sequences of cAR1 (uniprot id: P13773, residues: 1-268), cAR2 (uniprot id: P34907, residues: 1-267), cAR3 (uniprot id: P35352, residues: 1-260), P34907 and cAR4 (uniprot id: Q9TX43, residues: 1-269) were aligned using Clustal Omega (https://www.ebi.ac.uk/Tools/msa/clustalo/) in its default setting. Important residues known for class B are shown in red whilst the positional equivalents of those for cARs are shown in blue.

**Figure S4.**
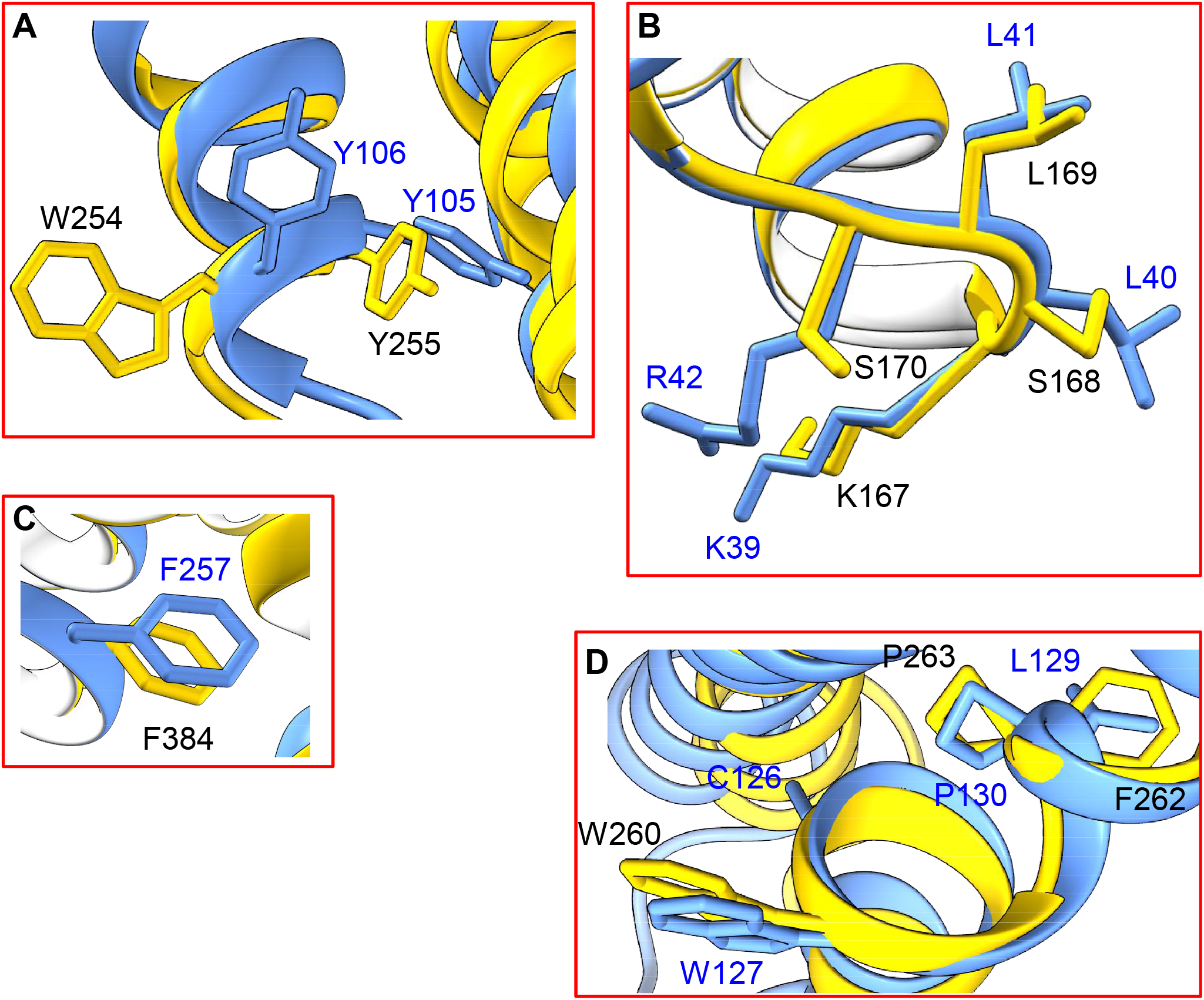
cAR residues that are potential positional equivalents of known GPCR motifs important for receptor structure and activation. cAR1 model A (shown in light blue) was superimposed on human CGRP receptor (shown in yellow, pdb: 6E3Y) using UCSF Chimera. Comparable residues are shown as sticks. Panel **A** shows the ^254^WY^255^ motif for hCGRP receptor and its positional equivalent, ^105^YY^106^ for cAR1. Panel **B** shows the positional equivalent of KKLH (or similar) motif within the ICL1 region that is shared between class A and B family of GPCRs. Panel **C** shows the positional equivalent of Y^7.53^ of the NPxxY^7.53^ motif in hCGRP receptor and cAR1. Panel **D** shows the GWGxP motif and its positional equivalent for hCGRP receptor and cAR1, respectively.

